# The neonatal Fc receptor is a pan-echovirus receptor

**DOI:** 10.1101/438358

**Authors:** Stefanie Morosky, Azia Evans, Kathryn Lemon, Sandra Schmus, Christopher J. Bakkenist, Carolyn B Coyne

**Affiliations:** Department of Pediatrics, University of Pittsburgh School of Medicine, Pittsburgh, PA USA; Center for Microbial Pathogenesis, UPMC Children’s Hospital of Pittsburgh, Pittsburgh, PA USA; Department of Radiation Oncology, University of Pittsburgh School of Medicine, Pittsburgh, PA USA; Department of Pharmacology and Chemical Biology, University of Pittsburgh School of Medicine, Pittsburgh, PA USA; R. K. Mellon Institute for Pediatric Research, UPMC Children’s Hospital of Pittsburgh, Pittsburgh, PA USA

## Abstract

Echoviruses are the main causative agents of aseptic meningitis worldwide and are particularly devastating in the neonatal population, where they are associated with severe hepatitis, neurological disease including meningitis and encephalitis, and even death. Here, we identify the neonatal Fc receptor (FcRn) as a pan-echovirus receptor. We show that loss of expression of FcRn or its binding partner beta 2 microglubulin (β2M) renders human brain microvascular cells resistant to infection by a panel of echoviruses at the stage of virus attachment and that a blocking antibody to β2M inhibit echovirus infection in cell lines and in primary human fetal intestinal epithelial cells. We also show that expression of human, but not mouse, FcRn renders non-permissive human and mouse cells sensitive to echovirus infection and that the extracellular domain of human FcRn directly binds echoviral particles and neutralizes infection. Lastly, we show that primary cells isolated from mice that express human FcRn are highly susceptible to echovirus infection. Our findings thus identify FcRn as a pan-echovirus receptor, which may explain the enhanced susceptibility of neonates to echovirus infections.

**Significance:** Echoviruses are associated with aseptic meningitis and induce severe disease, and even death, in neonates and young infants. Here, we identify the neonatal Fc receptor (FcRn) as a pan-echovirus receptor. FcRn is expressed on the surface of the human placenta, and throughout life in intestinal enterocytes, liver hepatocytes, and in the microvascular endothelial cells that line the blood-brain barrier. This pattern of expression is consistent with the organ sites targeted by echoviruses in humans, with the primary entry site of infection in the intestinal tract and subsequent infection of secondary tissues including the liver and brain. These findings provide important insights into echovirus pathogenesis and may explain the enhanced susceptibility of infants and neonates to echovirus-induced disease.

## Introduction

Echoviruses are small (~30nm) single stranded RNA viruses belonging to the *Picornaviridae* family. These viruses make up the largest subgroup of the Enterovirus genus and consist of approximately 30 serotypes. Enteroviruses are the main causative agents of aseptic meningitis worldwide, with echovirus 9 (E9) and echovirus 30 (E30) amongst the most commonly circulating serotypes (1). The neonatal and infant populations are at greatest risk for developing severe echovirus-induced disease and infection within the first few weeks of life can be fatal (2, 3). Enteroviral infections are also devastating in Neonatal Intensive Care Units (NICUs), where they account for 15-30% of NICU-associated nosocomial viral infections and result in death of the neonate in as many as 25% of cases (4–7), the majority of which result from echovirus 11 (E11) infections (8). In neonates, vertical transmission may occur at the time of delivery following a maternal infection in the days or weeks prior to delivery (9). In addition, echovirus infections have also been observed *in utero,* both at late and earlier stages of pregnancy, where they are associated with fetal death (10–14).

Echoviruses are primarily transmitted through the fecal-oral route where they target the gastrointestinal epithelium. In primary human fetal-derived enteroids, echoviruses exhibit a cell type specificity of infection and preferentially infect enterocytes (15). The basis for this cell-type specific tropism is unclear. Decay accelerating factor (DAF/CD55) functions as an attachment factor for some echoviruses (16), but DAF expression does not sensitize non-permissive cells to infection (17), suggesting that another cell surface molecule functions as the primary receptor. While integrin VLA-2 (α_2_β_1_) is a primary receptor for E1 (18), it does not serve as a receptor for other echoviruses. Other work has implicated a role for MHC class I receptors in echovirus infections due to inhibition of viral binding, entry, or infection by monoclonal antibodies to MHC class I and/or beta-2 microglobulin (β2M) (17, 19, 20), which is required for efficient cell surface trafficking of MHC class I receptors. However, the precise role for MHC class I and β2M remains unclear and the primary receptor for many echoviruses is unknown.

Here, we identify the human neonatal Fc receptor (FcRn) as a primary echovirus receptor. We show that human cells deficient in FcRn expression are resistant to echovirus infection and infection is restored by FcRn expression. Concomitantly, expression of human FcRn renders murine-derived cell lines and primary cells permissive to echovirus infection. In contrast, expression of the murine homolog of FcRn has no effect on viral infection in either human or mouse cells, identifying a species-specific role for FcRn in echovirus infection. Using primary human intestinal epithelial cell monolayers isolated from mid-gestation fetal small intestines, we show that a monoclonal antibody recognizing β2M, which non-covalently associates with FcRn and is required for FcRn cell surface expression (21), significantly reduces echovirus infection. Lastly, we show that recombinant FcRn in complex with β2M neutralizes echovirus infection and directly interacts with viral particles. Our data thus identify FcRn as a primary receptor for echoviruses, which has important implications for echovirus pathogenesis.

## Results

### Human cells deficient in FcRn are non-permissive to echovirus infection

We screened a panel of cell lines for their susceptibility to echovirus infection and found that human placental choriocarcinoma JEG-3 cells were resistant to infection by a panel of 7 echoviruses (E5-7, E9, E11, E13, and E30) but were highly permissive to the related enterovirus coxsackievirus B3 (CVB) (**Figure 1A**). Levels of echovirus infection in JEG-3 cells were comparable to those observed in mouse embryonic fibroblasts (MEFs), which are highly resistant to echovirus infection, and were significantly less than those observed in permissive cell types including human intestinal Caco-2, HeLa, human brain microvascular endothelial cells (HBMEC), and human osteosarcoma U2OS cells (**Figure 1A, Supplemental Figure 1A-C**). The resistance of JEG-3 cells to echovirus infection occurred at the level of viral binding or entry as infection was restored when cells were transfected with infectious viral RNA (vRNA) (**Supplemental Figure 1D**).

**Figure 1.**
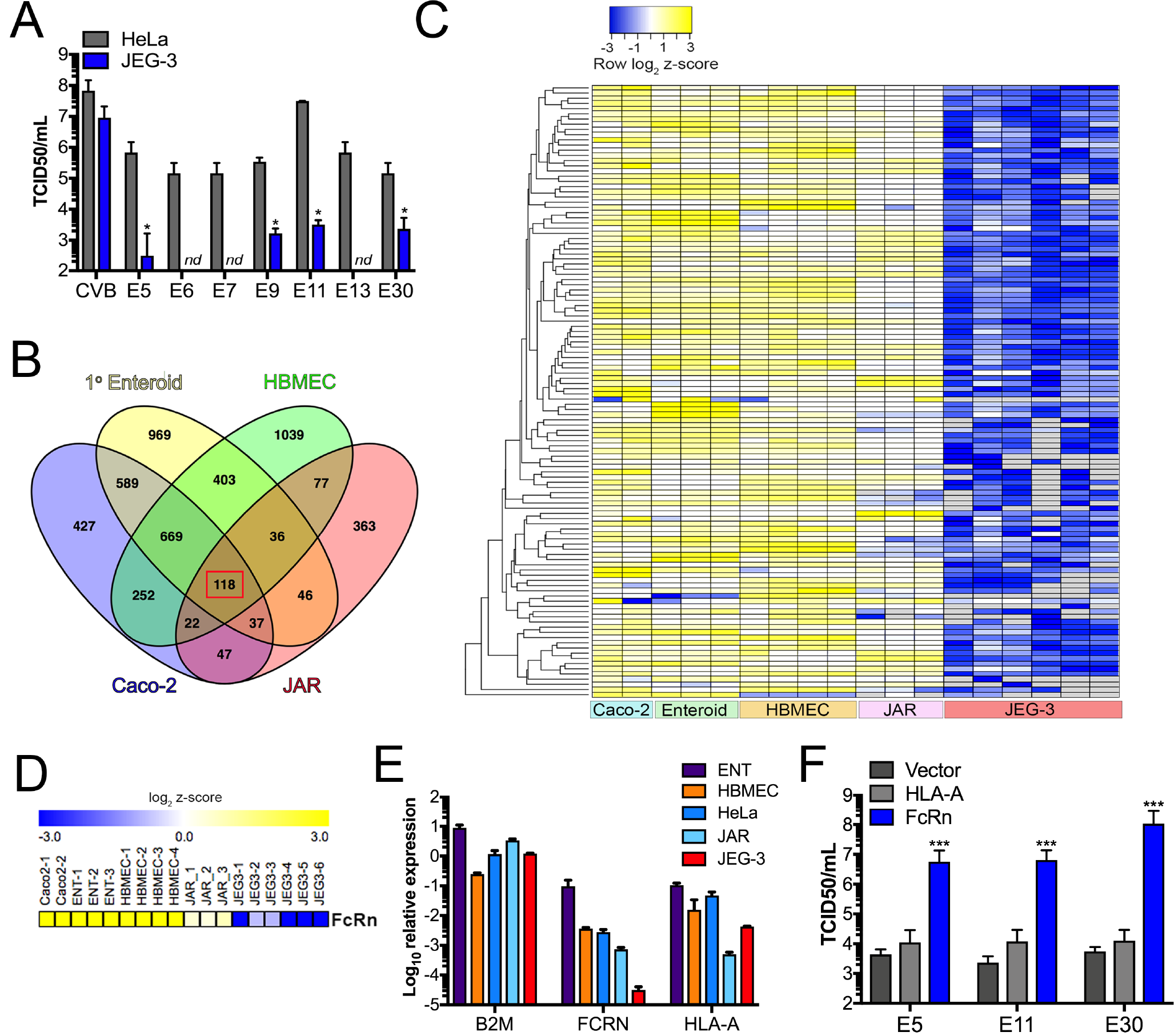
JEG-3 cells are not infected by echoviruses due to low FcRn expression. **(A)**, JEG-3 (blue bars) or HeLa (black bars) cells were infected with the indicated echovirus (1 PFU/cell) for ~24hrs. Viral titers (log_10_ TCID50/mL) from the indicated cell types are shown as mean ± standard deviation. Significance was determined using a t-test (p<0.05). **(B)**, Venn diagram from differential expression analysis using the DeSeq2 package in R between JEG-3 cells are either primary human fetal-derived enteroids (yellow), HBMEC (green), Caco-2 cells (blue), or JAR cells (red). There were 118 shared genes differentially downregulated between JEG-3 cells and these cell types (red square). **(C)**, Heatmap of 118 genes differentially downregulated in JEG-3 cells and the indicated cell type at bottom) based on log_2_ RPKM values. Transcripts with no reads are in grey. **(D)**, Heatmap of FcRn expression in the indicated cell type (based on log_2_ RPKM values). (E), RT-qPCR profiling of the level of expression of β2M (B2M), FcRn, or HLA-A in the indicated cell type. Data are shown as log_10_ relative expression normalized to actin and are shown as mean ± standard deviation. **(F)**, JEG-3 cells were transfected with vector control (pcDNA) or human HLA-A or FcRn for 24hrs, and then infected with the indicated echovirus for 24hrs. Viral titers (log_10_ TCID50/mL) are shown as mean ± standard deviation with significance determined with a Kruskal-Wallis test with Dunn’s test for multiple comparisons (***p<0.001). The relative expression of HLA-A and FcRn is shown in Supplemental Figure 1H.

We next performed RNAseq-based transcriptomics analyses between non-permissive JEG-3 cells and permissive cell types including Caco-2 and HBMEC cells and primary human enteroids harvested from fetal small intestines, which are highly sensitive to echovirus infection (15), to identify cell surface receptors differentially downregulated in JEG-3 cells. Because JEG-3 cells arise from choriocarcinomas and express many placental-specific transcripts, we also included JAR cells in our analyses, another human choriocarcinoma line that is more permissive to echovirus infection than JEG-3 cells (**Supplemental Figure 1E**). Using this approach, we identified 118 transcripts differentially downregulated in JEG-3 cells (p<0.001, log_2_ z score<-2, **Figures 1B, 1C** and **Supplemental Table 1**). Of these 118 transcripts, the neonatal Fc receptor (FCGRT, hereafter referred to as FcRn), was the most significantly downregulated cell surface receptor in JEG-3 cells (p<0.001, log_2_ z-score<-2), (**Figure 1D, Supplemental Table 1, Supplemental Figure 1F**). We confirmed the significantly lower levels of expression of FcRn in JEG-3 cells relative to permissive cell lines (HBMEC, HeLa, and JAR) and primary human fetal enteroids using RT-qPCR (**Figure 1D**). In contrast, there were no differences in expression of β2M, which is required to traffic FcRn to the cell surface (21) (**Figure 1E**). In addition, we confirmed previous findings that JAR cells are deficient in MHC class I molecules and that JEG-3 cells express very low levels of MHC class I molecules (22) (**Supplemental Figure 1G**), supporting the notion that these molecules were not responsible for the differential susceptibility of JEG-3 cells to echovirus infections.

To determine if the lack of FcRn expression was directly responsible for the low levels of echovirus infection in JEG-3 cells, we ectopically expressed human FcRn (hFcRn). Expression of hFcRn in JEG-3 cells significantly increased their susceptibility to infection by E5, E11, and E30 (~10,000-fold, **Figure 1F, Supplemental Figure 1H**). In contrast, expression of the related MHC class I or MHC class I-like molecules HLA-A and HLA-C and hemochromatosis protein (HFE), which also require β2M for cell surface expression, also had no effect on infection (**Figure 1F, Supplemental Figure H-I**). These data show that expression of hFcRn restores echovirus infection in non-permissive human cells.

### Expression of human FcRn restores echovirus infection in mouse cells

Echoviruses do not infect mouse cells efficiently (**Supplemental Figure 1A, 1B**). Since ectopic expression of hFcRn in human cells in which endogenous levels were low restored their susceptibility to infection, we next determined whether the murine homolog of FcRn (mFcRn) was also sufficient to promote infection. Whereas expression of hFcRn in JEG-3 cells restored infection of a panel of echoviruses (E5, E7, E11, E13, and E30) by ~10,000-fold, expression of mFcRn had no significant effect (**Figure 2A, Supplemental Figure 2A, 2B**). Similarly, we found that expression of hFcRn, but not mFcRn, rendered mouse embryonic fibroblasts (MEFs) and chinese hamster ovary (CHO) cells highly susceptible to echovirus infection (**Figure 2B, Supplemental Figure 2D**).

**Figure 2.**
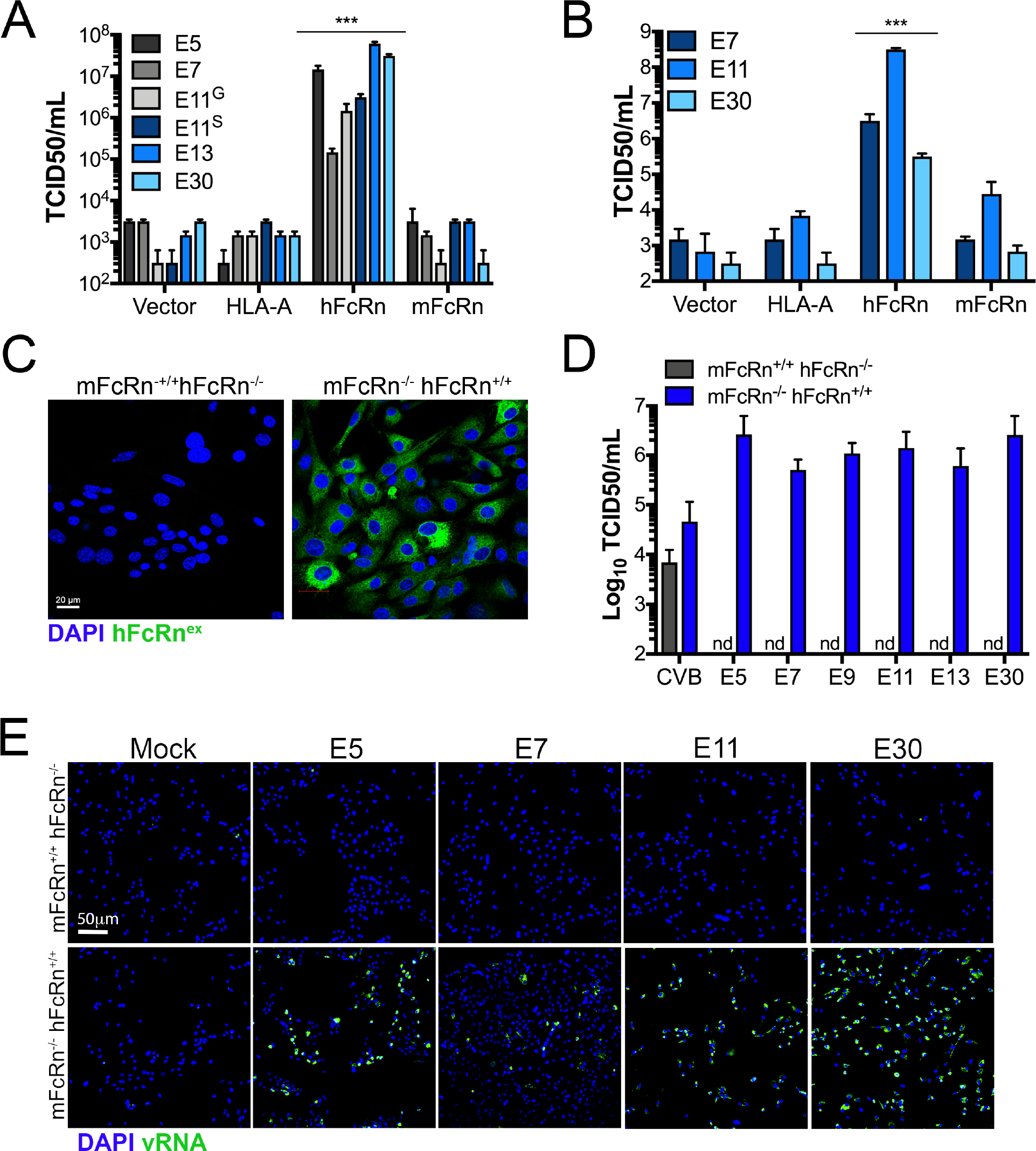
Human, but not mouse, FcRn expression restores infection in human-and mouse-derived cells. **(A)**, JEG-3 cells were transfected with vector control (pcDNA), human HLA-A, human FcRn (hFcRn), or mouse FcRn (mFcRn) for 24hrs and then infected with the indicated echovirus for 24h (E11^G^ Gregory strain and E11^S^ Silva strain). Viral titers (log_10_ TCID50/mL) are shown as mean ± standard deviation with significance determined with a Kruskal-Wallis test with Dunn’s test for multiple comparisons (p<0.001). The relative expression of HLA-A, hFcRn, and mFcRn is shown in Supplemental Figure 2B. **(B)**, Mouse embryonic fibroblasts (MEFs) were transfected with vector control (pcDNA), human HLA-A, or human or mouse FcRn (hFcRn, mFcRn, respectively) for 24hrs and then infected with the indicated echovirus for 24h. Viral titers (log_10_ TCID50/mL) are shown as mean ± standard deviation with significance determined with a Kruskal-Wallis test with Dunn’s test for multiple comparisons (p<0.001). The relative expression of HLA-A, hFcRn, and mFcRn is shown in Supplemental Figure 2C. **(C)**, Primary fibroblasts were isolated from mice expressing mouse, but not human, FcRn (mFcRn^+/+^ hFcRn^−/−^) or expressing human, but not mouse, FcRn (mFcRn^−/−^ hFcRn^+/+^) and then immunostained with an antibody recognizing the extracellular domain of hFcRn (in green). DAPI-stained nuclei are shown in blue. **(D)**, Primary fibroblasts isolated from mFcRn^+/+^ hFcRn^−/−^ mice (grey bars) or mFcRn^−/−^ hFcRn^+/+^ mice (blue bars) were infected with the indicated echovirus, or with coxsackievirus B (CVB) as a control for 24hrs. Viral titers (log_10_ TCID50/mL) are shown as mean ± standard deviation from cells isolated from four mice of each type. Nd, not detected. **(E)**, Primary fibroblasts isolated from mFcRn^+/+^ hFcRn^−/−^ or mFcRn^−/−^ hFcRn^+/+^ mice were infected with the indicated echovirus, or mock infected as a control, and then the level of viral replication assessed at 6hrs post-infection by immunofluorescence microscopy for double-stranded viral RNA (a replication intermediate, in green). DAPI-stained nuclei in blue. Scale bars are shown at bottom left in (C) and (E).

To further define the role of hFcRn in echovirus infection, we isolated primary fibroblasts from mice lacking expression of mFcRn, but expressing the α chain of hFcRn under the control of the endogenous promoter (mFcRn^−/−^ hFcRn^+/+^) or matched wild-type controls (mFcRn^+/+^ hFcRn^−/−^) (**Figure 2C**) (23, 24). In these cells, hFcRn is expressed at the cell surface in complex with mouse β2M. Primary fibroblasts isolated from mFcRn^+/+^ hFcRn^−/−^ mice were resistant to echovirus infection, as expected (**Figure 2D, 2E**). In contrast, cells isolated from mFcRn^−/−^ hFcRn^+/+^ mice were highly permissive to echovirus infection and exhibited >10,000-fold enhanced susceptibility to infection (**Figure 2D, 2E**). Collectively, these data show that expression of human, but not mouse FcRn is sufficient to confer cellular susceptibility to echovirus infection, which supports a species-specific role for FcRn in echovirus infections.

### Loss of FcRn expression renders cells resistant to echovirus infection

We next determined whether loss of FcRn expression rendered cells expressing FcRn less susceptible to infection. For these studies, we used RNAi-mediated silencing of FcRn in HBMEC, an immortalized human blood-brain barrier cell line that expresses high levels of FcRn (**Figure 1C, 1D**) and which is highly sensitive to echovirus infection (**Supplemental Figure 1A, 1B**). We found that silencing of FcRn expression by two independent siRNAs led to significant (~1000-10,000-fold) decreases in echovirus infection but had no effect on CVB infection (**Figure 3A, 3B, Supplemental Figure 3A**). Similar results were obtained in human osteosarcoma U2OS cells (**Supplemental Figure 3B**). In addition, silencing of β2M expression led to comparable reductions in infection (**Figure 3A-B, Supplemental Figure 3A-C**). In contrast, RNAi-mediated silencing of other cell surface molecules that require β2M for trafficking, such as HLA-A, HLA-B, HLA-C, and HFE had no significant effect on echovirus infection in HBMEC (**Supplemental Figure 3C**). Importantly, echovirus replication in β2M- and hFcRn-RNAi transfected cells was restored when cells were transfected with infectious vRNA (**Figure 3C**), suggesting that the inhibition occurred at the stage of virus binding or entry.

**Figure 3.**
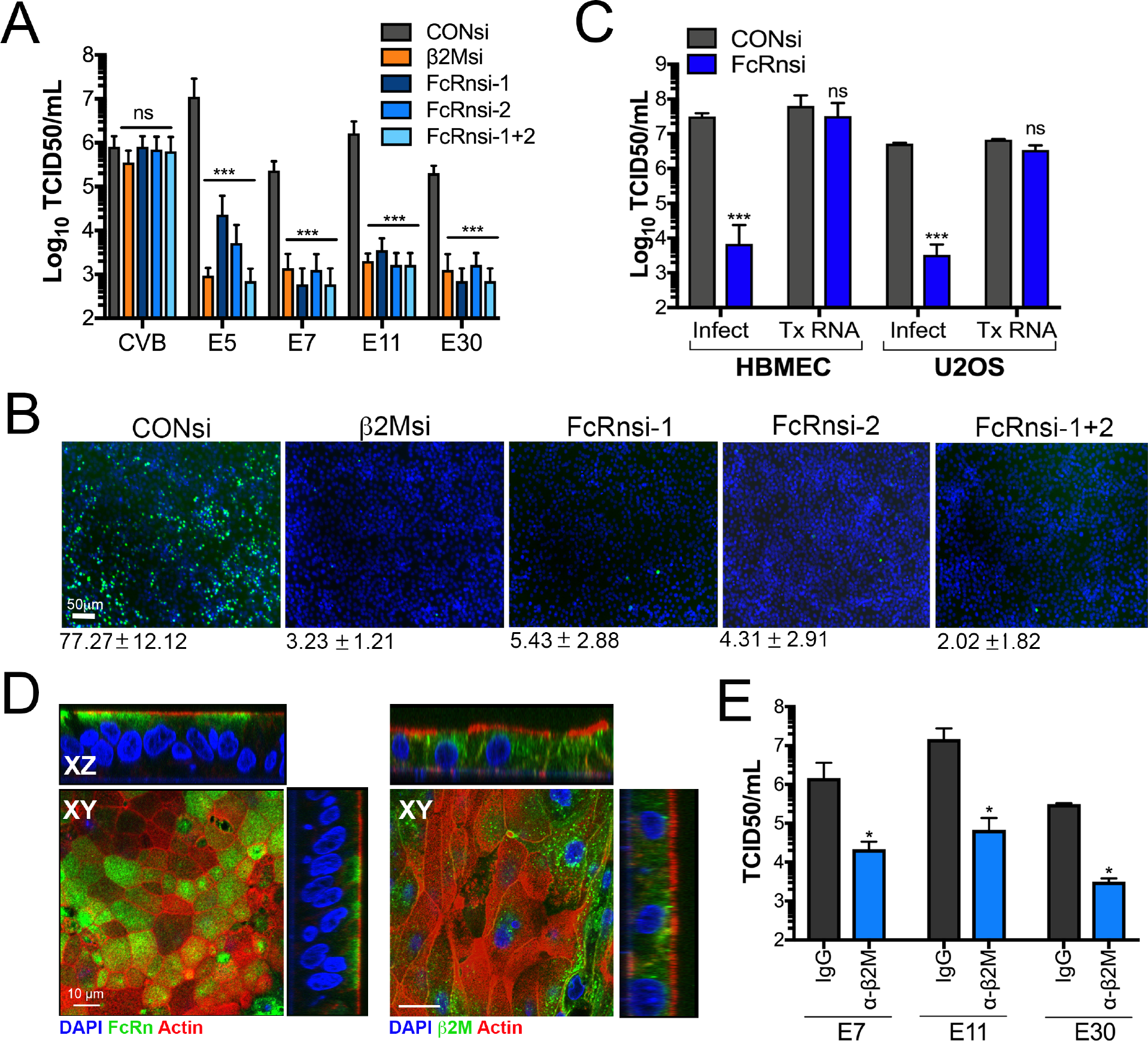
Loss of FcRn expression reduces echovirus infection. **(A)**, HBMEC were transfected with an siRNA against β2M (orange bar) or two independent siRNAs against FcRn (FcRn-1 and FcRn-2) alone or in combination (FcRn 1+2) (blue bars), or scrambled control siRNA (CONsi, grey bars) for 48hrs and then infected with CVB or the indicated echovirus for an additional 16hrs. Shown are viral titers (log10 TCID50/mL) as mean ± standard deviation with significance determined with a Kruskal-Wallis test with Dunn’s test for multiple comparisons (***p<0.001, ns not significant). **(B)**, HBMEC transfected with siRNAs as described in (A) were infected with E11 for 24hrs and then infection assessed by immunofluorescence microscopy for double stranded viral RNA (a replication intermediate, in green). DAPI-stained nuclei are shown in blue. The average level of infection (as determined by the percent of DAPI-stained nuclei that were positive for vRNA is shown at bottom as mean ± standard deviation. **(C)**, HBMEC transfected with scrambled control siRNA (CONsi) or FcRn siRNA (FcRnsi-1) for 48hrs were infected with E11 or transfected with infectious E11 viral RNA for an additional 24hrs. Shown are viral titers (log_10_ TCID50/mL) as mean ± standard deviation with significance determined with a t-test (***p<0.001, ns not significant). **(D)**, Confocal micrographs of fetal-derived primary human intestinal epithelial (HIE) cells immunostained for FcRn (left, green) or b2M (right, green) and counterstained for actin (left and right, red). DAPI-stained nuclei are shown in blue. Three-dimensional cross-sections are shown at top and right. Scale bars at bottom left. **(E)**, Primary fetal HIE were incubated with anti-β2M monoclonal antibody (blue bars) or isotype control antibody (grey bars) (2μg/mL for both) for 30min prior to infection with the indicated echovirus in the presence of antibody for an additional 24hrs. Shown are viral titers (log10 TCID50/mL) as mean ± standard deviation from three independent HIE preparations with significance determined with a t-test (*p<0.05).

To determine whether other echoviruses also required expression of FcRn, we performed a high content imaging-based screen using β2M siRNA and two siRNAs targeting FcRn (alone and in combination) for a panel of echoviruses (E5, E6-7, E9, E11, E13, E25, E29, and E30-32) as well as CVB as a control. We found that infection by all echoviruses tested was significantly reduced in cells depleted of β2M or FcRn expression while CVB infection was unchanged (**Supplemental Figure 3E**). In addition, consistent with previous studies that blocking antibodies to β2M inhibited infection of E7 and E11 (17, 19), we found that infection by E5, E7, E9, E11, E13, and E30 were also inhibited by a monoclonal antibody against β2M (**Supplemental Figure 3F**).

Because echoviruses are transmitted by the enteral route and infect the gastrointestinal epithelium as the primary site of host entry, we next determined whether FcRn was also involved in echovirus infection of intestinal epithelial cells. We showed previously that E11 preferentially infects enterocytes in human fetal-derived enteroids (15). To determine the role of FcRn in echovirus infection of the human neonatal intestine, we isolated intestinal crypts from human fetal small intestines (16–23w of gestation) and plated these crypts directly onto transwell inserts which leads to the formation of a fully differentiated single cell monolayer. We found that FcRn localized to the sub-apical domain of fetal-derived primary human intestinal epithelial (HIE) monolayers (**Figure 3D, left**), which is consistent with what has been observed *in vivo* in other polarized cells types, where FcRn localizes sub-apically and to the basolateral surface (25–29) whereas β2M was localized to both the basolateral contact sites and to intracellular vesicles (**Figure 3D, right**). Because primary HIE cannot be genetically altered, we used β2M monocloncal antibody to determine whether FcRn also facilitates echovirus infection of these cells. In HIE collected from three different human fetal intestine preparations, we found that β2M monoclonal antibody significantly reduced infection by E7, E11, and E30 (**Figure 3E**), similar to that which we observed in cell lines. Collectively, these data show that FcRn expression is required for echovirus infection and implicates a role for FcRn in mediating echovirus infection of the human neonatal intestine.

### FcRn facilitates echovirus attachment and directly interacts with viral particles

We found that echovirus infection in cells depleted of FcRn could be restored by transfection of cells with vRNA, which suggested that this inhibition occurred at the stage of viral binding or entry. We therefore determined whether downregulation of FcRn expression would alter echovirus binding. We found that silencing of FcRn expression in HBMEC significantly reduced cell surface binding of E5, E7, E9, E11, and E30 to HBMEC (**Figure 4A**). In contrast, this silencing had no effect on CVB binding (**Figure 4A**). Residual levels of viral binding in HBMEC may be mediated by cell surface factors such as DAF that facilitate binding of some echoviruses (16). Consistent with a role in viral binding, we also found that echovirus binding to primary mouse fibroblasts (mFcRn^−/−^, hFcRn^+/+^) was significantly higher than in cells expressing mFcRn (mFcRn^+/+^, hFcRn^−/−^) (**Figure 4B**). These data show that FcRn facilitates echovirus cell surface attachment.

**Figure 4.**
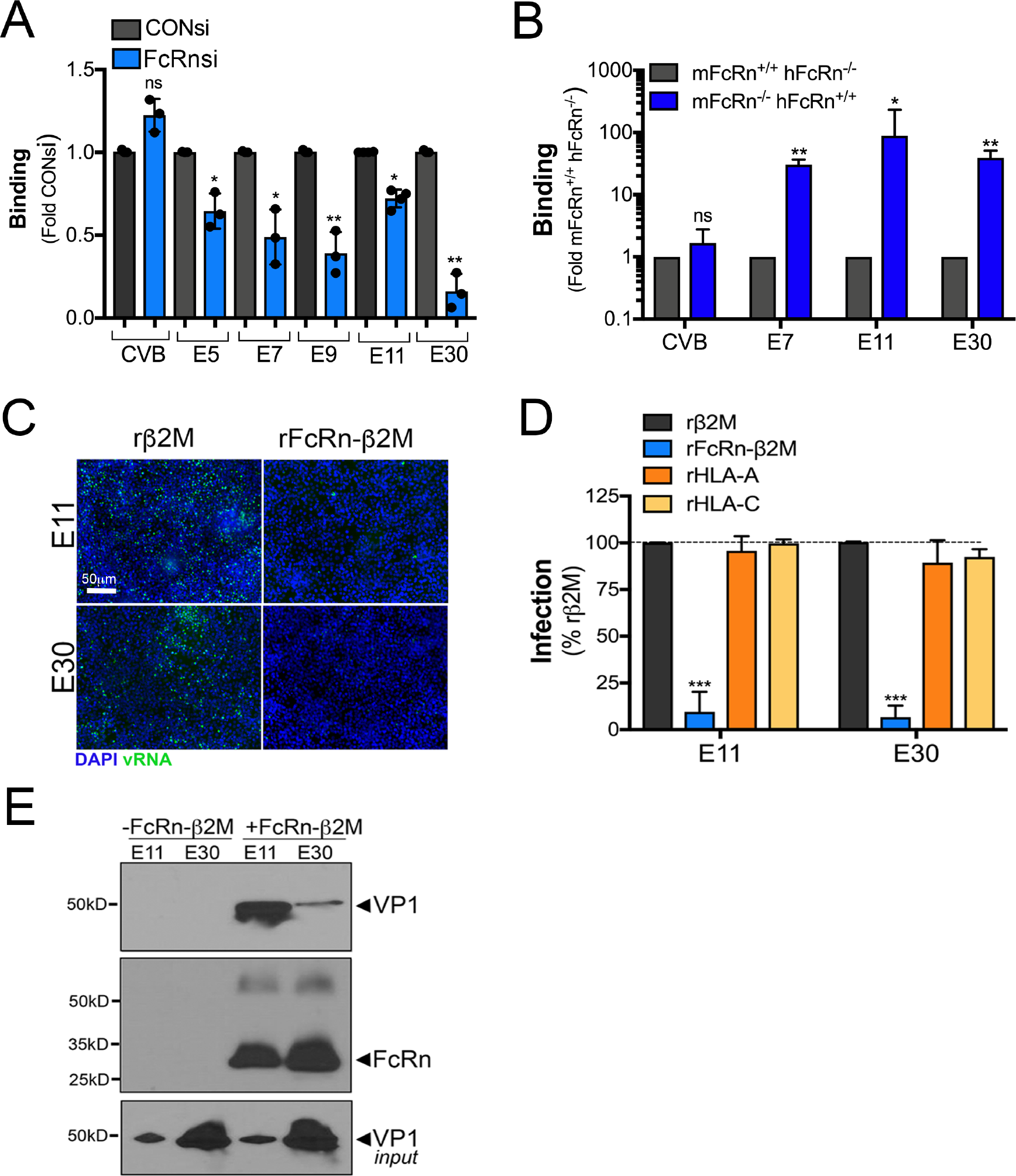
FcRn mediates echovirus binding. **(A)**, HBMEC were transfected with an siRNA against FcRn (FcRn-1, blue bars) or scrambled control siRNA (CONsi, grey bars) for 48hrs and then the extent of viral binding of CVB or the indicate echovirus (50 PFU/cell) as assessed by a RT-qPCR based binding assay. The extent of binding is shown as a fold from CONsi control (mean ± standard deviation). Significance was determined using a t-test (*p<0.05, ns not significant). **(B)**, The extent of viral binding of CVB or the indicated echovirus (50 PFU/cell) was assessed in primary fibroblasts isolated from mFcRn^+/+^ hFcRn^−/−^ or mFcRn^−/−^ hFcRn^+/+^ mice using a RT-qPCR based binding assay. Shown is the extent of binding in cells isolated from four mice of each type, which is shown as mean ± standard deviation. Significance was determined using a t-test (*p<0.05, **p<0.01, ns not significant). **(C)**, E11 or E30 (10^6^ particles) were incubated with recombinant β2M (rβ2M, 2.5μg/mL) or the extracellular domain of FcRn in complex with β2M (rFcRn-β2M, 2.5μg/mL) for 1hr at 4°C, pre-adsorbed to HBMEC for 1hr at 4°C, washed, and then cells infected for 16hrs. Shown are representative immunofluorescence images for double stranded viral RNA (a replication intermediate, green). DAPI-stained nuclei are shown in blue. **(D)**, E11 or E30 (10^6^ particles) were incubated with recombinant β2M (rβ2M, 2.5μg/mL), FcRn in complex with β2M (rFcRn-β2M, 2.5μg/mL), HLA-A (2.5μg/mL), or HLA-C (2.5μg/mL) for 1hr at 4°C, pre-adsorbed to HBMEC for 1hr at 4°C, washed, and then cells infected for 16hrs. The level of infection was assessed by immunostaining for vRNA normalized to DAPI-stained nuclei. Shown is the perfect of infection normalized to rB2M controls from experiments performed in triplicate (>1000 cells total) as mean ± standard deviation. Significance was determined with a Kruskal-Wallis test with Dunn’s test for multiple comparisons (***p<0.001). **(E)**, E11 or E30 (10^8^ particles) were incubated with recombinant β2M (rβ2M, 5μg/mL) or 6x His-tagged extracellular domain of FcRn in complex with β2M (rFcRn-β2M, 5μg/mL) for 1hr at 4°C, then incubated with Ni-NTA Agarose beads for 1hr at 4°C. Following extensive washing, immunoblots were performed for the viral capsid protein VP1 (top) and then membranes were stripped and re-probed with an antibody recognizing the extracellular domain of FcRn (middle). In parallel, level of input virus was immunoblotted with anti-VP1 antibody (bottom).

To determine whether FcRn directly interacts with echovirus particles, we used a recombinant protein approach utilizing a purified heterodimer containing the extracellular domain of FcRn in complex with β2M (rFcRn-β2M). We found that incubation of viral particles with rFcRn-β2M prior to infection neutralized both E11 and E30 infection (**Figure 4C, 4D**). In contrast, incubation with purified β2M alone, or recombinant HLA-A or HLA-C had no effect (**Figure 4C, 4D**). To determine whether there was a direct interaction between FcRn and echoviral particles, we performed *in vitro* binding assays using rFcRn-β2M. Using this approach, we found that rFcRn-β2M co-precipitated with purified E11 and E30 in *in vitro* binding assays, (**Figure 4E**), demonstrating a direct interaction between FcRn and echovirus particles.

## Discussion

Here, we identify FcRn as a primary receptor for echoviruses. We show that expression of FcRn is necessary and sufficient for echovirus infection and that FcRn directly binds echovirus particles and facilitates viral binding. We also show that expression of human, but not mouse FcRn restores echovirus infection in non-permissive mouse and human cells and thereby identify a species-specific mechanism of infection. Our data show that a number of clinically relevant echoviruses commonly associated with human disease, including E9, E30, and E11, utilize FcRn as a receptor, suggesting a pan-echovirus role. In contrast, FcRn plays no role in the infection of related enteroviruses including CVB and PV. Our findings provide important insights into the cellular receptor used by echoviruses to initiate their infections, which also provides further insights into a variety if aspects of echovirus pathogenesis.

FcRn transports and regulates the circulating half-life of IgG throughout life (30–32). In addition, FcRn is responsible for the development of passive immunity through the transfer of maternal-derived antibodies. In humans, expression of FcRn on the placenta (33) is solely responsible for the establishment of passive immunity in the fetus due to transport of maternal-derived IgG across the placental surface directly into fetal blood (34). This differs in rodents, where passive immunity is established postnatally from maternal-derived IgG in milk/colostrum (35). FcRn is expressed throughout life in a variety of cell types including the small intestine (26, 36), the microvasculature of the blood-brain barrier (37), myeloid cells (38), hepatocytes (39, 40), amongst others. Although echoviruses are primarily transmitted via the fecal-oral route, viral-induced disease is associated with infection of secondary organs, most notably the liver and brain. The expression of FcRn on the surfaces of the intestine, brain microvasculature, and heptocytes may thus explain the tropism of echoviruses for these tissues and the viral mechanism to bypass the barriers presented by the cells comprising these sites.

FcRn binds albumin as well as IgG (39, 41). Although FcRn binds to albumin and IgG at distinct sites (42), both of these interactions occur within the low pH (≤6.5) environment of endosomes, with release occurring in the basic pH (≤7.5) of the bloodstream. In contrast, our findings reveal a direct interaction between echoviruses and FcRn that occurs at the neutral pH of the cell surface prior to viral entry. Once internalized, it is possible that the interaction between FcRn and echoviruses is altered by the low pH of endosomes, which may facilitate subsequent genome release and/or endosomal escape. We have shown that E11 preferentially infects enterocytes (15), with enhanced infection from the basolateral surface of HIE (43). This polarity of infection is consistent with the enhanced expression of FcRn in enterocytes in the intestine and its enrichment to the basolateral surface. Following replication, E11 is released bidirectionally from HIE from both the apical and basolateral domains (43). Given that FcRn mediates bidirectional transport (29), this raises the possibility that echoviruses could be transported from either the apical or basolateral domains to cross the intestinal barrier.

Echoviruses are associated with severe disease in neonates, particularly during the first two weeks of life and in those born prematurely. The vertical transmission of echoviruses is thought to occur at the time of delivery and be associated with maternal infection in the preceding days or weeks. However, fetal infections *in utero* have also been associated with disease and/or death (10–14), suggesting that vertical transmission might also occur during pregnancy. FcRn is highly expressed on syncytiotrophoblasts (30, 44), the fetal-derived cells that comprise the outermost cellular barrier of the human placenta that directly contact maternal blood. These cells are highly resistant to viral infections due to intrinsic antiviral defense pathways (33). However, given that FcRn expressed on the surface of these cells transcytoses maternal-derived IgG directly into the underlying fetal blood. Our identification of FcRn as an echovirus receptor raises the possibility that echoviruses might have higher rates of transplacental transfer than has been appreciated. In addition, it should be noted that the highest rates of transplacental IgG transfer occur in the third trimester, with the level of maternal-derived IgG is greater in the fetus than in the mother (45). Thus, a maternal echovirus infection in the later stages of pregnancy could potentially lead to FcRn-mediated placental infection or transplacental viral transport and expose the fetus to virus prior to delivery. Further defining the role of FcRn in echovirus infections *in utero* and postnatally will provide important insights into echovirus-induced fetal and neonatal disease.

Our work presented here identifies FcRn as a pan-echovirus receptor. Given that FcRn-based therapeutics have been developed to target a variety of human diseases (46), our findings also point to FcRn as a possible target for anti-echovirus therapeutics to ameliorate virus-induced disease. Future studies identifying the mechanism by which echoviruses utilize FcRn to enter or bypass barrier tissues such as the GI epithelium, blood-brain barrier, and placenta will provide important insights into a variety of aspects of echovirus pathogenesis.

## Materials and Methods

### Additional Materials and Methods are located in Supplemental Information

#### Cell lines

Human brain microvascular cells (HBMEC) were obtained from Kwang Sik Kim (Johns Hopkins University) and have been described previously (47) and were grown in RPMI-1640 supplemented with 10% fetal bovine serum (FBS, Invitrogen), 10% NuSerum (Corning), non-essential amino acids (Invitrogen), sodium pyruvate, MEM vitamin solution (Invitrogen), and penicillin/streptomycin. JEG-3, JAR, U2OS, and Caco-2 (BBE clone) cells were purchased from the ATCC and were cultured as described previously (48, 49). HeLa cells (clone 7B) were provided by Jeffrey Bergelson (Children’s Hospital of Philadelphia) and were cultured in MEM supplemented with 5% FBS and penicillin/streptomycin.

#### Primary cells

Experimental procedures were approved by the University of Pittsburgh Animal Care and Use Committee and all methods were performed in accordance with the relevant guidelines and regulations. Primary fibroblasts were generated from 4 week old B6.Cg-Fcgrt<tm1Dcr> Tg(CAG-FCGRT) 276Dcr/DcrJ (Cat. 004919) and control C57BL/6J (000664) mice purchased from The Jackson Laboratory (Bar Harbor, Maine). Mice were euthanized according to institution standards and ears and tail were removed, incubated in 70% ethanol for 5min and then rinsed twice in PBS + 50ug/mL kanamycin for 5 min. Hair was removed and tissue was cut into small pieces and incubated in 9.4mg/mL collagenase D (Roche, 11088858001) and 1.2 mg/mL pronase (Roche, 1088858001) in complete DMEM at 37°C with shaking at 200rpm for 90min. The resulting cell suspensions were filtered through 70uM cell strainers, collected at 580g, resuspended in complete DMEM containing 10 units penicillin and 10ug streptomycin/mL and 250ng/mL amphotericin B and cultured at 37°C in a humidified 5% CO2 incubator.

Primary human intestinal epithelial cells were isolated from crypts isolated from human fetal small intestines as described (15). Complete methods can be found in Supplemental Materials and Methods.

#### Viruses and viral infections

Experiments were performed with coxsackievirus B3 (CVB) RD strain, poliovirus (PV, sabin strain, type 2), echovirus 5 (Noyce stain, E5), echovirus 6 (Burgess strain, E6), echovirus 7 (Wallace strain, E7), echovirus 9 (Hill strain, E9), echovirus 11 (Gregory or Silva strains, E11^G^ and E11^S^), echovirus 13 (Del Carmen, E13), echovirus 25 (JV-4, E25), echovirus 29 (JV-10, E29), echovirus 30 (Bastianni strain, E30), echovirus 31 (Caldwell strain, E31), or echovirus 32 (PR-10 strain, E32) that were provided by Jeffrey Bergelson (Children’s Hospital of Philadelphia) and were originally obtained from the ATCC. Viruses were propagated in HeLa cells and purified by ultracentrifugation over a sucrose cushion, as described (50).

Unless otherwise stated, infections were performed with 1 PFU/cell of the indicated virus. In some cases, viruses were pre-adsorbed to cells for 1hr at 4°C in serum-free MEM supplemented with 10mM HEPES followed by extensive washing in 1x PBS or complete media. Infections were then initiated by shifting cells to 37°C for the times indicated. Viral titers were determined by TCID50 assays in HeLa cells using crystal violet staining.

Binding assays were performed by pre-adsorbing 50 PFU/cell of the indicated virus to cells for 1hr at 4°C in serum-free MEM supplemented with 10mM HEPES followed by extensive washing with 1x PBS. Immediately following washing, RNA was isolated, and RT-qPCR performed for viral genome-specific primers, as described below.

For experiments using blocking antibodies, cells were incubated with the indicated antibodies (at 5μg/mL) for 1hr at 4°C in serum-free DMEM containing 10mM HEPES. For anti-DAF IF7 blocking experiments, all incubations were performed in DMEM containing 10% FBS and 10mM HEPES. Following this incubation, viruses were pre-adsorbed to cells in the presence of antibodies for an additional 1hr at 4oC in serum-free or serum-containing medium, washed extensively, and then cells infected at 37oC for the indicated time in the presence of

#### Plasmids, siRNAs, and transfections

Sequence verified vectors (pcDNA 3.1) expressing human HLA-A (, HLA-C, FcRn or mouse FcRn were purchased from Genscript. EGFP-fused HFE (pCB6-HFE-EGFP) was a gift from Pamela Bjorkman (Addgene plasmid # 12104) and was described previously (51). Plasmids were reverse (MEFs, CHO cells) or forward (JEG-3 cells) transfected with the indicated plasmids using Lipofectamine 3000 according to the manufacturer’s instructions.

Pooled siRNAs (four total) targeting HLA-A and HFE were purchased from Dharmacon (siGENOME, M-012850-01 and M-011051-02). Pooled siRNAs (four total) targeting HLA-B and HLA-C were purchased from Santa Cruz Biotechnology (sc-42922 and sc-105525). Control (scrambled) siRNA was purchased from Sigma (Mission Universal, SIC001). Individual siRNAs targeting were synthesized by Sigma, with sequences as follows: (β2M UCCAUCCGACAUUGAAGUU; FcRn-1 CCACAGAUCUGAGGAUCAA; FcRn-2 ACUUUUGACUGUUAGUGAC). In all cell types, siRNAs were reverse transfected into cells using Dharmafect-1 (Dharmacon) according to the manufacturer’s instructions.

#### Antibodies

The following antibodies or reagents were used—recombinant anti-dsRNA antibody (provided by Abraham Brass, University of Massachusetts and described previously (52)), mouse monoclonal anti-VP1 (NCL-ENTERO, Leica), mouse monoclonal anti-FcRn (Santa Cruz Biotechnology, sc-271745), rabbit polyclonal FcRn (Abcam ab139152), rabbit monoclonal HLA-A (Abcam, ab52922), rabbit monoclonal HLA-C (Abcam, ab126722), PE-conjugated anti-HLA antibody (recognizing HLA A-C, HLA-E) (Novus, NBP2-68006PE), mouse monoclonal anti-β2M (Sigma, SAB4700010), and isotype control mouse monoclonal IgG antibody (MOPC 21, Sigma, M5284). Alexa-fluor 594 conjugated phalloidin was purchased from Invitrogen (A12381). Anti-DAF IF7 antibody was provided by Jeffrey Bergelson (Children’s Hospital of Philadelphia).

#### Recombinant protein, *in vitro* pulldowns, and immunoblotting

Purified native β2M was purchased from Bio-rad (6240-0824) and was isolated from human urine. Recombinant HLA-A and HLA-C was purchased from Novus (H00003105 and NBP2-2310, respectively). Recombinant extracellular domain of FcRn in complex with β2M was purchased from Sino Biological (CT009-H08H) and was purified from HEK293 cells. For viral neutralization studies, purified viral particles (10^6^) were incubated with the indicated recombinant protein (2μg) for 1hr at 4°C with constant rotation in serum-free MEM supplemented with 10mM HEPES. This complex was then added to cells for an additional 1hr at 4°C, cells washed extensively with 1x PBS, and infections initiated by shifting to 37oC for the 16-24hrs, as indicated in figure legends.

*In vitro* pulldowns between E11 and E30 were performed by incubating purified virus particles (10^7^) with 2μg of purified 6xHis tagged FcRn complex to β2M for 1hr at 4°C with constant rotation in buffer containing 100mM NaCl, 20mM Tris-Cl (pH 7.4), 0.5mM EDTA, and 0.5% (v/v Nonidet-40). Following this incubation, HiPur Ni-NTA agarose beads were added for an additional 1hr at 4oC with constant rotation. Bead complexes were then pelleted by centrifugation and washed 6x with wash buffer (10mM Tris-Cl (pH 8.0), 1mM EDTA, 1% Triton X-100, 0.1% SDS, and 140mM NaCl). Beads were then resuspended in denaturing sample buffer and immunoblots performed, as described below.

For immunoblotting, the lysates described above were loaded onto 4-20% Tris-HCl gels (Bio-Rad) and transferred to nitrocellulose membranes. Membranes were blocked in 5% nonfat dry milk, probed with the indicated antibodies, and developed with horseradish peroxidase-conjugated secondary antibodies (Santa Cruz Biotechnology), and SuperSignal West Dura chemiluminescent substrates (Pierce Biotechnology). Membranes were stripped for reprobing using ReBlot Strong antibody stripping solution (Millipore, 2504) according to the manufacturer’s instructions.

#### Immunofluorescence microscopy

Cells were washed with PBS and fixed with ice-cold 100% methanol for immunostaining of viral infections or with 4% paraformaldehyde at room temperature, followed by 0.25% Triton X-100 to permeabilize cell membranes for a minimum of 15min at room temperature for all other immunostaining. Cells were incubated with primary antibodies for 1 hour at room temperature, washed with 1x PBS, and then incubated for 30 minutes at room temperature with Alexa-Fluor-conjugated secondary antibodies (Invitrogen). Slides were washed and mounted with Vectashield (Vector Laboratories) containing 4′,6-diamidino-2-phenylindole (DAPI). Images were captured using a Zeiss LSM 710 inverted laser scanning confocal microscope or with inverted IX81 or IX83 Olympus fluorescent microscopes. Images were adjusted for brightness/contrast using Adobe Photoshop (Adobe). Image quantification for the extent of infection was performed using Fiji (Cell counter plugin) or the CellSens Count and Measure package, as indicated. A minimum of 1000 cells were quantified.

#### RT-qPCR

Total RNA was prepared using the Sigma GenElute total mammalian RNA miniprep kit, according to the protocol of the manufacturer. RNA was reverse transcribed with the iScript cDNA synthesis kit (Bio-Rad) following the manufacturer’s instructions. A total of 1 μg of total RNA was reversed transcribed in a 20 μL reaction, and subsequently diluted to 100 μL for use. RT-qPCR was performed using the iQ SYBR Green Supermix or iTaq Universal SYBR Green Supermix (Bio-Rad) on a CFX96 Touch Real-Time PCR Detection System (Bio-Rad). Gene expression was determined based on a ∆*C*_*Q*_ method, normalized to human actin. Primer sequences to actin, CVB, and pan-echovirus primers have been described previously (49, 53). Primers to β2M, FcRn, HLA-A, HLA-B, HLA-C, and HFE were synthesized by Sigma and sequences can be found in Supplemental Table 2.

#### Statistics

All statistical analysis was performed using GraphPad Prism. Experiments were performed at least three times. Data are presented as mean ± standard deviation. A Student’s t-test or One-Way Anova was used to determine statistical significance, as described in figure legends. P values of < 0.05 were considered statistically significant, with specific P-values noted in the figure legends.

## Acknowledgements

We thank Jacqueline Corry and Charles Good (UPMC Children’s Hospital of Pittsburgh for technical assistance, Jeffrey Bergelson (Children’s Hospital of Philadelphia) for helpful suggestions and reagents, Kwang Sik Kim (Johns Hopkins University) for providing HBMEC, and Terrence Dermody (UPMC Children’s Hospital of Pittsburgh) for helpful suggestions. This project was supported by NIH R01-AI081759 (C.B.C.), a Burroughs Wellcome Investigators in the Pathogenesis of Infectious Disease Award (C.B.C), and the Children’s Hospital of Pittsburgh of the UPMC Health System (C.B.C). This project also used the UPMC Hillman Cancer Center and Tissue and Research Pathology/Pitt Biospecimen Core and Animal Facility shared resources which are supported in part by award P30CA047904.

